# Immigration from the metacommunity affects bdelloid rotifer community dynamics most

**DOI:** 10.1101/450627

**Authors:** Nicolas Debortoli, Frederik De Laender, Karine Van Doninck

## Abstract

Dispersal is an important driver of local community dynamics. It has been proposed that, for communities composed of microscopic organisms, dispersal could well be the dominant process, outpacing local processes driven by environmental conditions and species interactions. This is because microscopic organisms often reproduce asexually, fostering rapid colonization, and are easily dispersed by water or air current. We studied the case of bdelloid rotifers belonging to the genus *Adineta*, microscopic asexual animals with dried stages that are easily dispersed by wind to investigate the relative effects of dispersal and local processes on their community dynamics. To this end, we constructed a classic competition model to theoretically examine how spatial and local biodiversity dynamics varied with fitness and dispersal characteristics of bdelloid *Adineta* species. Next, we compared our predictions with an experimental dataset containing spatio-temporal *Adineta* community dynamics from the wild. This comparison suggested that immigration from the local meta-community was the most critical parameter under the conditions tested. One *Adineta vaga* species, abundant in the surrounding area, rapidly colonized our experimental habitats and dominated most of the communities. We also ran the model under different levels of environmental conditions (permissive, intermediate and harsh) to simulate seasonal community variability and found that communities experience important bottlenecks yearly in winter but that the same community re-established. The dissimilarities observed between roof communities suggest differences in adaptation or immigration capacities. Besides their asexual reproduction and extreme desiccation tolerance, a key characteristic of bdelloid ecology identified here, is the spatio-temporal dynamic of abundant bdelloid clones present in the meta-community that rapidly colonize empty patches to establish new populations.

## Introduction

Dispersal is an essential driver of community dynamics, affecting species distributions and abundances. Micro-organisms (<2mm) mostly rely on passive dispersal through transport on larger animals or through wind erosion. This type of dispersal results in a more randomized immigration pattern, with possible long-range dispersals (Ptatscheck et al., 2018) and with global (gamma) diversity in some micro-organisms being similar to their local (alpha) diversity (Finlay, 2002). Some studies even suggest that the distribution of microscopic species is a function of habitat preferences rather than of dispersal limitation or historical contingency (Bass and Cavalier-Smith, 2004; Costello and Chaudhary, 2017). This has also been suggested for bdelloid rotifers, a clade of microscopic animals (<2mm) showing distinct specificities for habitat type and strong dispersal capacity. Bdelloid rotifers are very abundant micro-organisms in semi-terrestrial habitats such as soil, moss and lichen patches (Fontaneto *et al*, 2006a; Fontaneto *et al*, 2007; Fontaneto *et al*, 2008) forming small propagules, called “tuns”, when entering a dormant stage during long periods of drought or freezing (Ricci, 1998; Ricci and Caprioli, 2005). This “tun” formation, which can occur at any stage of their life cycle, provides bdelloid individuals with strong dispersal capacity, being easily blown by the wind or attached to microscopic sediments that can passively be transported over large distances, enabling a large geographic distribution (Fontaneto and Ricci, 2006b; Wilson, 2011). Available data however also reveal patchy local distribution of bdelloids, with almost no overlap in species composition among different substratum (Fontaneto *et al*, 2006a; Fontaneto *et al*, 2007; Fontaneto *et al*, 2008), suggesting specificity to habitat type (Fontaneto *et al*, 2006a; Fontaneto and Ricci, 2006b; Fontaneto *et al*, 2011).

While the life-span of bdelloid rotifers rarely exceeds 30 days, propagules can persist several years in desiccated state and eventually return to an active state upon rehydration (Guidetti and Jönsson, 2002). Survival after desiccation has never been quantified under harsh environmental conditions, but laboratory experiments demonstrated that survival could be extremely high under controlled permissive conditions depending on the protocol used (Ricci, 1998; Hespeels *et al*, 2014). Survival could reach 60-100% when groups of 10-20 individuals of *Adineta vaga or Philodina vorax* were desiccated on filter paper respectively (Ricci, 1998). When > 100 *A. vaga* individuals were placed on low melting point agarose, a desiccated cluster was formed and the survival rate reached > 80% after 84 days of desiccation (Hespeels *et al*, 2014). Ricci and Caprioli (2005) even described a reproductive boost following rehydration of the desiccated bdelloid propagules. Desiccation therefore appears a critical component of the bdelloid rotifers life-cycle, also providing a way out from parasites (Wilson *et al*, 2010; Wilson, 2011). It has indeed been shown that the association of bdelloid rotifers with their *Rotiferophthora* parasites could be disrupted when dry periods were coupled with wind dispersal, the parasites being less desiccation-tolerant.

Besides their extreme tolerance to desiccation, freezing and high doses of ionizing radiation, bdelloid rotifers are also remarkable for their asexual mode of reproduction (Maynard Smith, 1978). No males or meiosis have ever been observed in bdelloids and each female is able to clone herself, which allows rapid colonization of new habitats starting from a single individual (hydrated or dried).

While several studies have monitored the spatial distribution of bdelloid rotifer species, showing large dispersal capacities and low but significant habitat preferences (Fontaneto *et al*, 2006a and 2011), only one study included the temporal dynamics, describing no variation in species composition with seasonality (Ricci *et al*, 1989). This latter study however focused on a single large moss patch from which subsamples were collected monthly. It is therefore difficult to disentangle which ecological parameters (dispersal, habitat preferences, species interactions, environmental variations…) may impact the community structure of bdelloid rotifers, each subsample being interdependent. We may therefore wonder how bdelloid communities are spatially structured over time, knowing that these micro-animals reproduce asexually, disperse easily as tuns and resist harsh conditions.

Here, we studied the spatial and temporal dynamics of bdelloid species communities carrying out a controlled, quantitative field experiment during two years. Every three months we sampled nine communities (present in petri-dishes) located on three distinct roofs in Namur (Belgium) and isolated and genotyped each bdelloid *Adineta* individual. We then analyzed the data using a competition model. Using the model, we simulated colonization and the subsequent community dynamics of the *Adineta* individuals based on solely three main parameters: the immigration rate (i.e. the dispersal capacities of each species), the survival probability, and the reproductive output of each individual with the two latter parameters being indicative of species fitness.

We ran the model under five distinct scenarios (Figure 1), making various assumptions on how fitness and dispersal contributed to the local dynamics, and compared the resulting simulations to our experimental data on *Adineta*. In the first scenario, we assumed all species have similar fitness (Adineta species have similar feeding capacities as they all graze on the substrate surface), similar dispersal rate (size of ‘tuns’ is comparable) and having closely related life-cycles resembling the assumptions underlying neutral models (Hubell, 2001). Species co-occurrence is only short-lived and due to stochastic events (birth-death-immigration). In a second scenario, similar dispersal rates were also considered, but distinct habitat preferences depicted by distinct probabilities to survive and reproduce (Fontaneto *et al*, 2006a and 2011). In a third scenario, the dispersal rate varied across bdelloid species but not the fitness. We define dispersal rate as the chance for one individual to be passively transported to the community from the metacommunity in which extremely abundant species, at a given place, could immigrate more frequently in nearby habitat patches (Amarasekare and Nisbet, 2001; Lowe and McPeek, 2014). In a fourth scenario, fitness and dispersal rate covaried negatively. This scenario reflects the observation of Ricci and Caprioli (2005) that life-history traits of bdelloid rotifers could vary with habitat preferences, providing evidence for a shorter life span, a higher fecundity and earlier age at first reproduction in bdelloid species that do not desiccate and inhabit permanent freshwater bodies. As a consequence, bdelloid species less tolerant to desiccation should have limited dispersal capacities but a higher reproductive output. Finally, in a fifth scenario all parameters covaried positively. We postulated that one or a few species are widely represented in the nearby area, favoring the chances for short-range dispersal and persistence. We included this scenario because Debortoli *et al* (2016) observed that the two most abundant species in the local community (named species A and C) were also the one present in most samples (39-44% of the lichens sampled) (see also Ricci et al, 1989). It has been shown in other microscopic species that community similarity decreased significantly with geographical distance, more than with environmental distance (Soininen et al, 2007).

**Figure 1:**
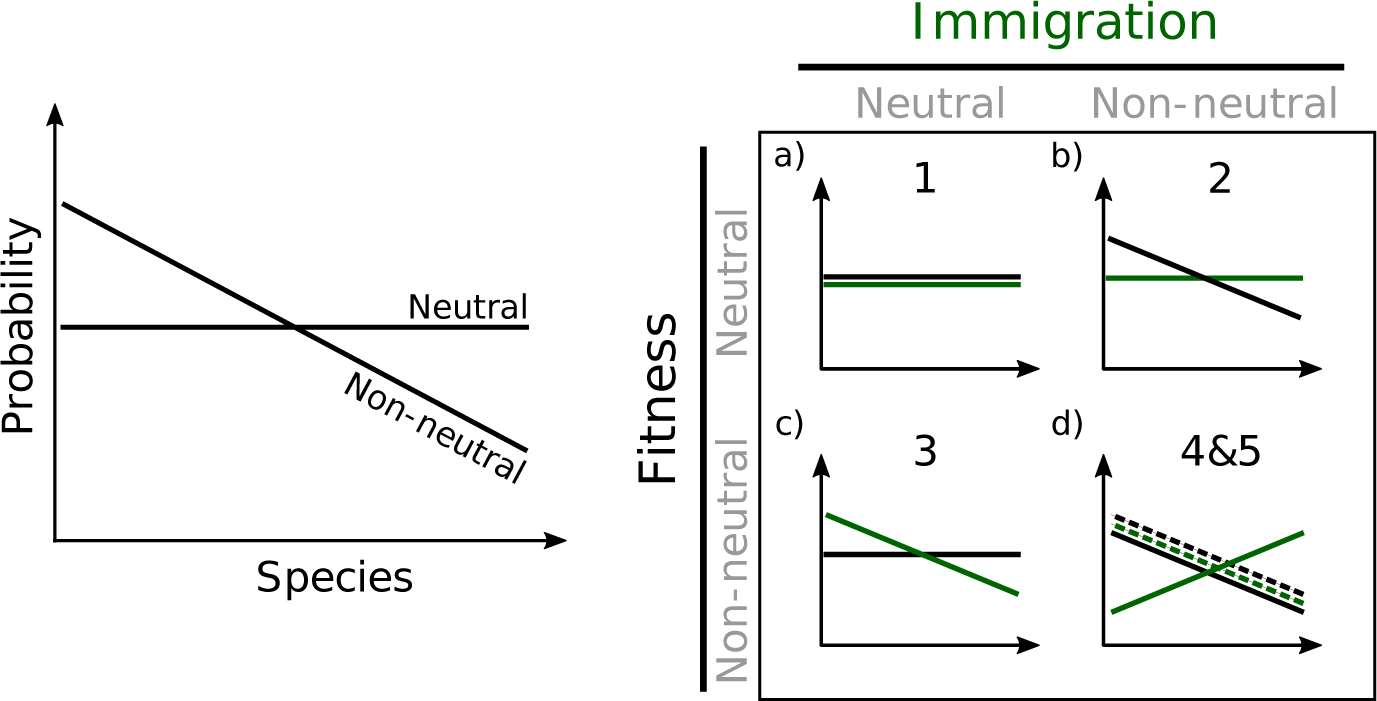
Probability distribution tested for the model under five distinct scenarios. On the left panel: two scenarios are presented. In one scenario, the probabilities of given parameters are identical across all species (neutral). In the other scenario, the probabilities vary across species (non-neutral). On the right panel: five scenarios build from distinct combinations (neutral and non-neutral) of two parameters (the species-specific probability to immigrate in the community (in green) and fitness (in black)) are shown. a) In scenario 1, all species have the same probabilities to immigrate, reproduce and survive at each time-step (linear distribution). b) In scenario 2, the probability to immigrate is identical across species, but the probability distributions to reproduce and survive vary. c) Scenario 3 simulates conditions opposite to scenario 2, reproduction and survival are equal for all species but the probability to immigrate varies across species. d) Solid lines represent scenario 4 modelling a trade-off between the probabilities to immigrate and the probabilities to reproduce and survive. The dashed lines on d) represent scenario 5 in which probabilities to immigrate, reproduce and survive at each time-step co-vary.

## Materials and methods

### Sampling and species delimitation

We selected three flat roofs located across the UNamur campus (Belgium, Namur; 50°27′58.27”N; 4°51′37.76”E) on each of which we placed three Petri dishes (Ø = 10 cm) filled with 20mL of 3% solid agarose. To avoid any contamination the dishes were kept closed with Parafilm^®^ until being placed on the respective roofs. The three Petri dishes from a same roof were located 8-25 meters from each other whereas 32-148 meters separated the dishes from distinct roofs. Each Petri dish was fixed on a block and maintained leaned for three months, being exposed to environmental conditions and to passive dispersal of micro-organisms (Figure 2a). This kind of setting resulted in a humid but not submerged community mimicking the limno-terrestrial habitat of bdelloid rotifers with the layer of agarose buffering humidity variations and forming a suitable substrate for bdelloid rotifers (Wilson and Sherman, 2013). Every three months, each dish on each roof was carefully sealed with Parafilm^®^ and brought back to the laboratory (Figure 2b) for morphological identification and isolation of each *Adineta* individual morphologically determined following Donner (1965). Our isolation protocol consisted in washing each bdelloid rotifer by pipetting it into successive clean Spa^®^ water drops and transferring it to an individual tube. Each tube was then briefly centrifuged to pellet the isolated animal and inspected under a binocular to make sure that only a single rotifer was present. If no individual was found in the tube, the tube was discarded (no second individual was added into the tube to avoid contamination).

**Figure 2:**
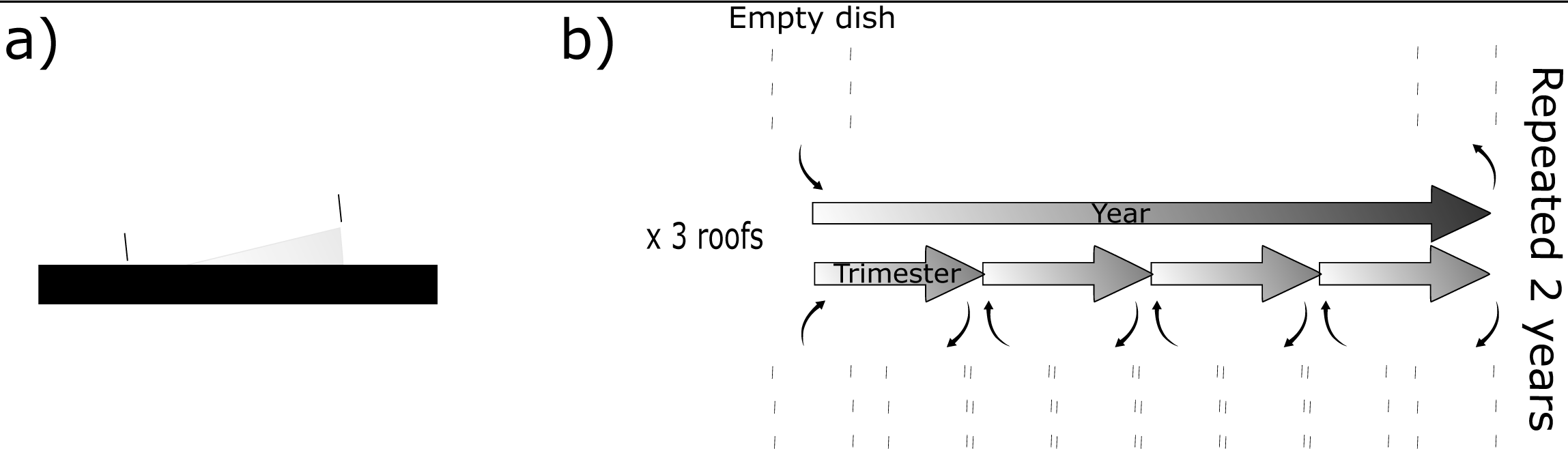
Experimental design for Adineta vaga sampling on the roofs of UNamur campus (Belgium, Namur; 50°27′58.27”N; 4°51′37.76”E). a) Representation of the freshly placed 10cm^2^ Petri dish, without lid, that was coated with 3% agarose [Sigma reference #9012-36-6]and maintained inclined to enable moisture without bathing. b) On each of the three roofs (A, B and C), three distinct empty petri-dishes were placed and replaced every three months with fresh dishes. The replaced dishes were brought back to the laboratory for community analyses. In parallel, three additional dishes were placed on each roof for a period of one full year before being replaced. This experiment was conducted for two years.

In parallel, we did place one Petri dish, prepared with the same method, next to each of the trimestral dishes and left it one full year on each roof. It was then collected and analyzed like the others. The experiment took place from December 2013 to December 2015, in total 8 samplings of the trimestral dishes were done and 2 samplings of the yearly dishes (Figure 2b).

The genomic DNA of each isolated *Adineta* individual was then extracted using the Chelex^®^ protocol: each sample was mixed with 35µL of Chelex^®^ solution [InstaGene^TM^ Matrix, Bio-Rad, #7326030] and 1 µL of proteinase K [Qiagen, #19133], homogenized by vortexing, heated 20 min at 56°C and 10 min at 95°C. Then, the Chelex^®^ beads were precipitated for 5 min at 14000 rpm and the supernatant containing the genomic DNA was transferred to a new tube and stored at -20°C. We used the data from Debortoli et al (2016) in which all individuals sampled were genotyped and calculated a rarefaction curve (‘vegan’ R package; Oksanen et al, 2007) to delimit the minimum subset of individuals to genotype in order to retrieve accurate species richness (Supplemental figure 1). As a result, for each combination of dish per sampling period, a subset of 24 individuals were randomly selected to amplify a portion of the mitochondrial cytochrome c oxidase subunit I (COI) gene using Folmer’s universal primers (HCOI: 5’ - TAA ACT TCA GGG TGA CCA AAA AAT CA - 3’ and LCOI: 5’ - GGT CAA CAA ATC ATA AAG ATA TTG G - 3’; Folmer *et al*, 1994). The PCR conditions were the same as in Debortoli *et al* (2016), except that the quantity of gDNA used was 5µL and that the number of cycles was set to 60. The amplicons were sent for Sanger sequencing to the GENEWIZ facilities (anciently Beckman-Coulter Genomics, UK).

The COI sequences obtained were aligned with 258 *Adineta sp*. sequences from GenBank and 378 *Adineta sp*. sequences from our own projects using MAFFT (E-INS-i method; Katoh and Standley, 2013) and visualized in MEGA5 (Tamura *et al*, 2011). One sequence was selected for each distinct haplotype resulting in a final dataset of 364 COI unique sequences (Supplemental table 1) of 589bp which was used for the ultrametric Bayesian tree reconstruction (BEAST v1.6.2; Drummond and Rambaut, 2007). We used the same parameters than previously applied to bdelloid rotifers to generate the trees (Tang *et al*, 2014; Debortoli *et al*, 2016) except that we combined 3 independent analyses with the LogCombiner package to avoid MCMC to be stuck in a local optimum. The resulting tree was used as input for species delimitation by the General-Mixed Yule Coalescent method with single threshold (Pons *et al*, 2006; Fujisawa and Barraclough, 2013).

### Community diversity across roofs and seasons

We represented our entire dataset (i.e. all COI genotyped *Adineta* individuals sampled on the roofs) on a species matrix clustered in order to group communities (i.e. each petri-dish) that present similar species assemblages using “heatmap3” package on R (Zhao *et al*, 2014). For each community at each timepoint where *Adineta* individuals were retrieved (81 in total), we calculated the total abundance (i.e. all *Adineta* individuals morphologically identified in the petri-dish), the number of *Adineta* species delimited by the GMYC method (referred to as “richness”), and Pielou’s Index of evenness (Pielou, 1966). We also calculated the Bray-Curtis indices of dissimilarities for each pair of communities using the “vegan” R package (Bray and Curtis, 1957; Oksanen *et al*, 2007). Then, we computed a Mantel test to identify whether dissimilarities between communities was related to geographic distance and ANOVA tests to identify which parameters (seasons and roofs) affected the different metrics calculated for each community (abundance, species richness, Pielou’s index of evenness, spatial dissimilarities). The meteorological data were provided by the Royal Meteorological Institute (IRM, see Supplemental table 2) which has a recording station located just outside Namur. Finally, we ran a factorial analysis (R, “ade4” package) to define species assemblages and community structures.

### Model

We used a discrete time competition model (Beverton and Holt, 1957; Hart et al, 2016), but allowing overlapping generations. The model incorporates three processes which intuitively apply to bdelloid rotifers: reproduction, passive immigration, and survival

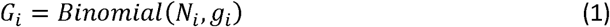

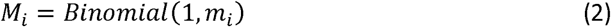

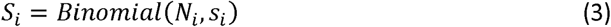

where N_i_ is the abundance of species i; g_i_ is its probability to reproduce; m _i_is its probability to immigrate in the community; and s_i_ is its probability to survive. G_i_ is the number of individuals from this species that will reproduce. M_i_ is the number of immigrating individuals of species i (either 0 or 1). S_i_ is the number of individuals of species i that survive.

The actual reproductive output from the reproducing individuals is given by *G*_*i*_ *C*_*i*_, where C_i_ measures the intensity of competition within and among species:

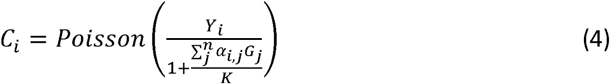

with Y_i_ the fecundity per individual of species i, K is the habitat size, and α_i, j_ is the intra- and inter-specific interaction strength.

Thus, the equation allowing intraspecific variations and the overall dynamics of species i is given by:

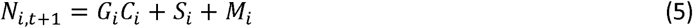

### Parameters setting

#### 1) Default parameters

To our knowledge, no studies measured survival or reproduction in field conditions for bdelloid species. Ricci et al (1983) observed that, in laboratory conditions (at 20°C), *Adineta vaga* lived 17 days on average with the main reproduction occurring during the early days of maturity and then slowly decreasing until death. They calculated that one *A. vaga* individual produced on average 1.2 eggs per day throughout its reproductive days, that the mean generation time was 7.7 days and that the intrinsic population increase was 0.344. As a result, we set Y equal to 1 for all species, corresponding to realistic numbers of eggs laid per individual at each time-step (*i.e.* one week or one generation) (Supplemental figure 2a). We also performed in our laboratory a life-cycle experiment with *A. vaga* AD008 clone and we observed one reproductive peak at day 6 during which fecundity reached 6 eggs, we estimated that this was due to controlled laboratory conditions and rarely the case in nature (see Supplemental data). We simplified the model by considering that fecundity was independent of the individual age and we implemented a stable fecundity throughout the simulations. We tested several combinations of values for Y and g_i_ and compared the population growth under those parameters to the rate of natural increase in *A. vaga* (Ricci, 1983). Setting Y=1 and g_i_=0.3 produced the best results as the population growth under those values was approximately 0.4 as in Ricci (1983) (Supplemental figure 2b). In addition, under those parameters the total number of eggs produced per capita was estimated to be 12 on average which fits the observation that *A. vaga* has a net reproduction output of 14.3 on average (Ricci, 1983) (Supplemental figure 2c). We always ran the model for 12 time-steps, each representing one week, to simulate three months of community dynamics, which corresponded to our field experiment. For the parameters defined here, the maximum number of individuals for the species with the highest reproductive output would be 60 individuals after 12 time-steps on average.

We tested several survival probabilities (s_i_) and obtained that 0.9 was the most accurate value in order to have around 40% of the initial population still alive after 12 time-steps, as observed in our preliminary studies on clone AD008 (Supplemental figure 2d and Supplemental data). The average number of immigrants per week was calculated in a preliminary study during which we used the same experimental design, but sampled plates weekly. There were *Adineta* individuals in 13 out of 48 Petri dishes (13/48 = 0.271) after one week and 20 *Adineta* individuals in total (20/48 =0.417). As three dishes presented more than one individual, we could not determine if each rotifer immigrated independently or if the first colonizer reproduced; we therefore decided to set the probability to immigrate from the metacommunity (m_i_) to 0.3 for *Adineta* species. Finally, we set the habitat size (K) to 150 as the maximum number of *Adineta* individuals found in a single lichen patch by Debortoli et al (2016) was 157.

The number of species present in the metacommunity described in the model was inferred from the GMYC analysis. With this approach, we considered all *Adineta* species referenced in public databases and in the present study. Because of their peculiar dispersal capacities, we postulate that all the *Adineta* species have a non-null probability to immigrate in our local samples, although variations may exists among species.

#### 2) Experimental factors

We ran the model according to the five scenarios explained in the introduction (Figure 1 and Supplemental figure 3). Furthermore, we considered that the indices of population growth calculated by Ricci et al (1983) in optimal laboratory conditions probably overestimated the reproductive output in harsher field conditions. We therefore ran each of the five scenarios three times: reproduction, survival and dispersal probabilities were 2 times lower in intermediate conditions (orange in Supplemental figure 3) and 10 times lower in harsh conditions (red) than in permissive ones (green).

We ran all simulations using four different types of probability distributions for g_i_, s_i_ and m_i_: linear, exponential and normal, as well as a hybrid between the exponential and normal distribution (Supplemental figure 3). Except for scenario 1, we predicate that probabilities to immigrate, reproduce and survive at each time-step are not equal across all species. The exponential and the normal distributions represent differences among species, but one (exponential) or a few (normal) species are supposed to be particularly efficient at immigrating and well adapted in comparison with the other species (Ricci *et al*, 1989; Fontaneto *et al*, 2011; Debortoli *et al*, 2016). We also tested a hybrid between an exponential and a normal distribution that would represent the case in which one species adapted to the local conditions outcompetes a few other species that are able to colonize and persist in sub-optimal conditions, e.g. species from alike habitats located in the nearby area. For scenario 1, only the linear distribution is applicable.

Finally, because, to our knowledge, no information about the ecological interactions between bdelloid species are available, we ran the model under three arbitrary intensities of intra- and inter-species interaction strength (with α_i, j_ = 0, 0.1 and 0.2).

### Comparison with field data

For each combination of scenario (5), conditions (3), distribution shape (4) and species interaction strength (3), we simulated 100 communities. We started all simulations with n species (delimited by the GMYC analysis) of null abundance to represent the empty communities at t_0_. At each of the 12 time-steps, the frequency of each species was recorded.

In order to estimate the accuracy of the model, we compared the simulations with the experimental data as the difference Δ between data and predictions (De Laender *et al*, 2014):

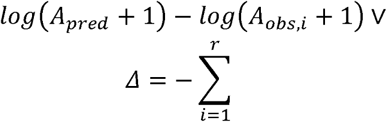

Where r is the number of communities and A is the variable of interest (the relative abundance of each species across the whole simulation, the abundance of individuals in each patch, species richness, evenness, or spatial dissimilarities) at t=12. Values of Δ approximating zero indicate a better model fit. Then, the sum of all the Δ calculated for each simulation was calculated to compare global best model fit. The model and all statistics were run in R, using the “vegan” package for Pielou’s index of species evenness and spatial β-diversity (Oksanen *et al*, 2007).

## Results

### Metacommunity structure

We isolated bdelloid individuals morphologically identified as *Adineta* from 81 out of the 90 roof patches sampled during two years, representing a total of 2663 *Adineta* individuals (median = 16.5 individuals per patch, range = 0-118). The sampling details and the subset of 1169 individuals for which we successfully sequenced the mtCOI marker are described in Supp. Table 3. This represented a total of 56 distinct mtCOI haplotypes, all corresponding to *Adineta vaga*, that were combined with all the published COI sequences from the genus *Adineta* to delimit genetic clusters (i.e. species) using the General-Mixed Yule coalescent approach. This method significantly delimited 117 species within *Adineta* (confidence interval: 111-135, number of ML entities identified being 119 containing 2 outgroup sequences; LR test: 0.0), 24 A. vaga species were retrieved in our two-year study on the roofs (Supplemental figure 4) confirming that this represents a cryptic species (Fontaneto et al, 2007; Fontaneto and Barraclough, 2015; Debortoli et al, 2016).

Interestingly, eight of the 24 species retrieved during our experiment (species 6, 7, 9, 13, 14, 15, 16, 19) have already been sampled from lichen patches throughout Belgium during the SPEEDY project (Merckx *et al*, 2018), four species (7, 9, 14 and 19) corresponded to species also previously detected in the park of Namur next to the University roofs (referred to as Species A, B, C and F in Debortoli *et al*, 2016) (Figure 3). The dominant species 9, also found throughout Belgium (Figure 3), represented 788 out of the 1169 individuals (67.41%) genotyped and was retrieved in 64 roof patches (71.11%) during the two years of this study (Figure 4). Two other species (species 8 and 13) were present in several roof patches sampled (22 and 18 patches respectively) but were much less abundant (93 and 97 individuals respectively) and not widespread within Belgium. The other species were less frequently found in our Petri dishes and were present at lower densities (see supplemental data for details and Figure 4). Interestingly, five species present on our roofs were also retrieved outside Belgium (Figure 3 and Supplemental figure 5).

**Figure 3:**
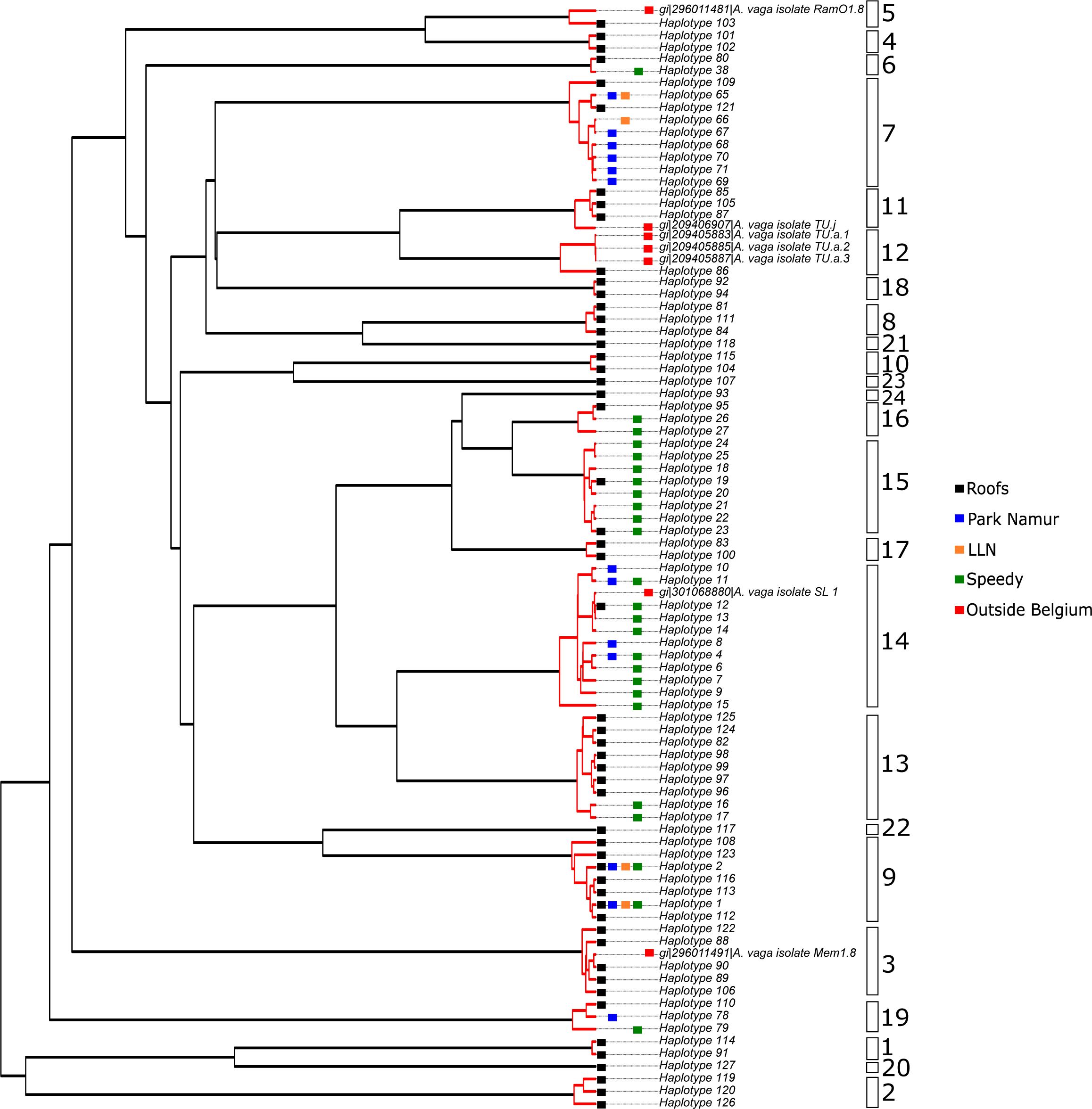
Pruned ultrametric tree built from our local mtCOI dataset and Adineta sp. COI sequences available on GenBank. Only the 24 GMYC-defined species that were collected on the roof experiment are represented (boxes) in this simplified version of the ultrametric tree (Supplemental figure 4 represents the entire dataset). Haplotypes that cluster within a same evolutionary entity are linked by red branches. Black squares indicate the haplotypes from the our experimental roof dataset whereas blue, orange, green and red squares highlight the haplotypes that were retrieved from other studies and were sampled in Namur, Louvain-La-Neuve (LLN), Flanders (Speedy project) and outside Belgium, respectively.

### Community diversity across roofs and seasons

The species richness in each roof patch ranged from 0 to 5 (median = 2) and Pielou’s index of species evenness ranged from 0 to 1 (median = 0.66). The linear models revealed that roof location (A, B and C) did not influence significantly neither abundance of individuals sampled nor species evenness and only slightly richness (p-value = 0.045, adjusted R^2^ = 0.047) with a lower diversity on roof C (Table 1, Figure 4). In contrast, these three metrics were significantly affected by seasons (Table 1). Indeed, the number of individuals sampled was significantly higher in autumn (median = 30, range 1-118) and lower in winter (median = 9.5, range = 0-60) than in the other seasons (medians = 22.5-26, ranges = 0-93) as suggested by the ANOVA (Table 1, Figure 5). We also calculated which meteorological parameters (T°, relative humidity, rainfall and wind) varied the most across seasons. Autumn was characterized by slightly higher minimum humidity than other seasons whereas summer and spring were much drier (df = 3; R^2^ = 0.424; p < 2e^-16^; Supplemental table 4, Supplemental figure 6). Temperatures were also significantly lower in winter than in other seasons (df = 3; R^2^ = 0.604; p < 2e^-16^, Supplemental figure 6). Rainfall and wind intensity varied significantly among seasons but those variations were small (df = 3; R^2^ = 0.015 and 0.056; p < 0.005 and 5.35e^-09^, respectively, Supplemental figure 6).

**Figure 4:**
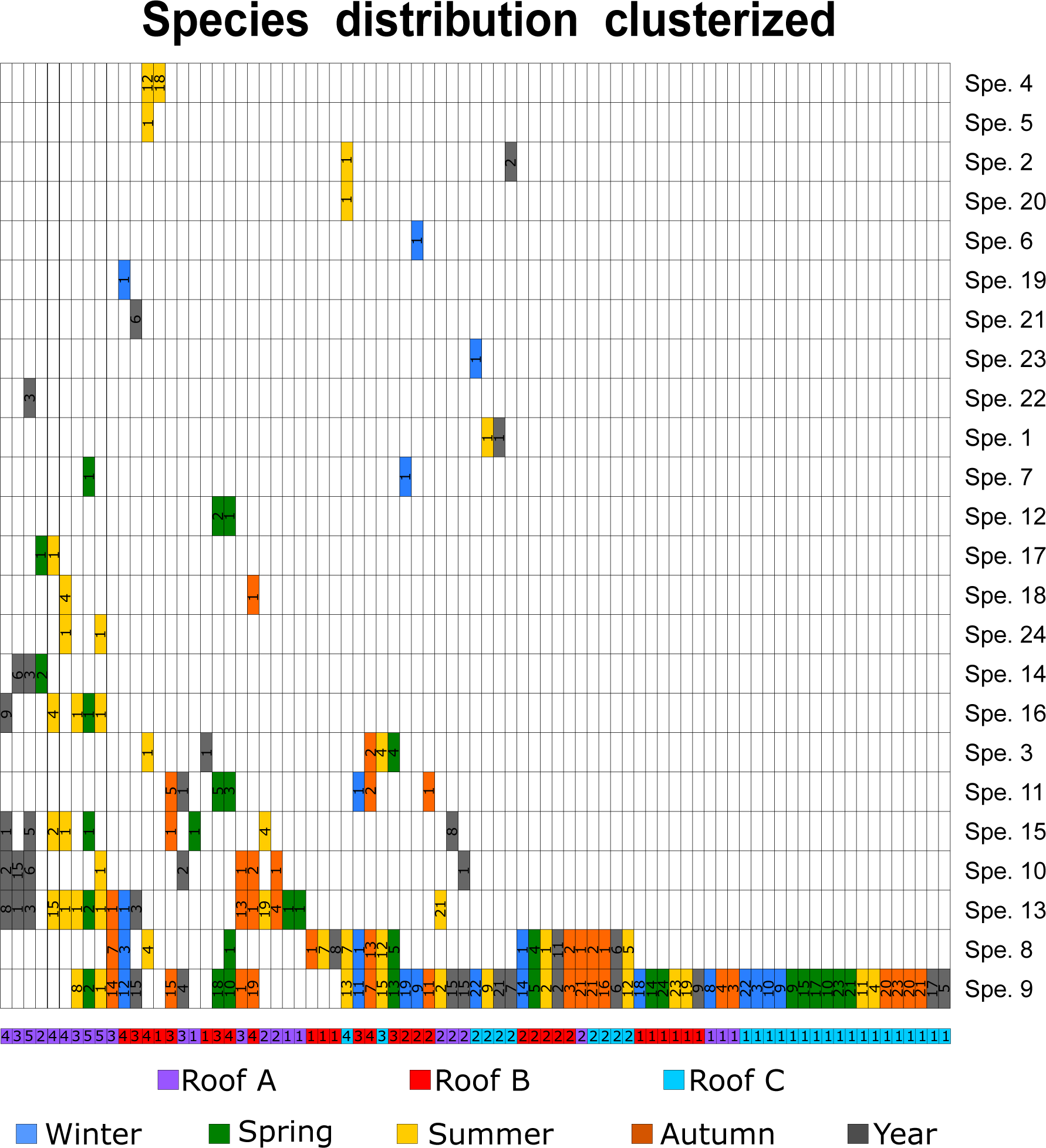
Clustered matrix of Adineta vaga species distribution. The matrix was built from the presence/absence of each A. vaga species (as defined by GMYC in Figure 3) in each non-empty community collected on the roofs (81 communities or petri-dishes), and the communities were then clustered hierarchically according to species composition. Colored boxes indicate the presence of a species in a specific dish or community. The box color represents the season at which this species was present and the boxes under the matrix show on which roof it was sampled (A, B and C are in violet, red and turquoise, respectively). Grey boxes indicate that the species was sampled in the dishes collected yearly. The number of individuals genotyped for each species in each patch is indicated in the boxes. The species richness found within each patch is indicated in the patch box, under the matrix.

**Table 1:**
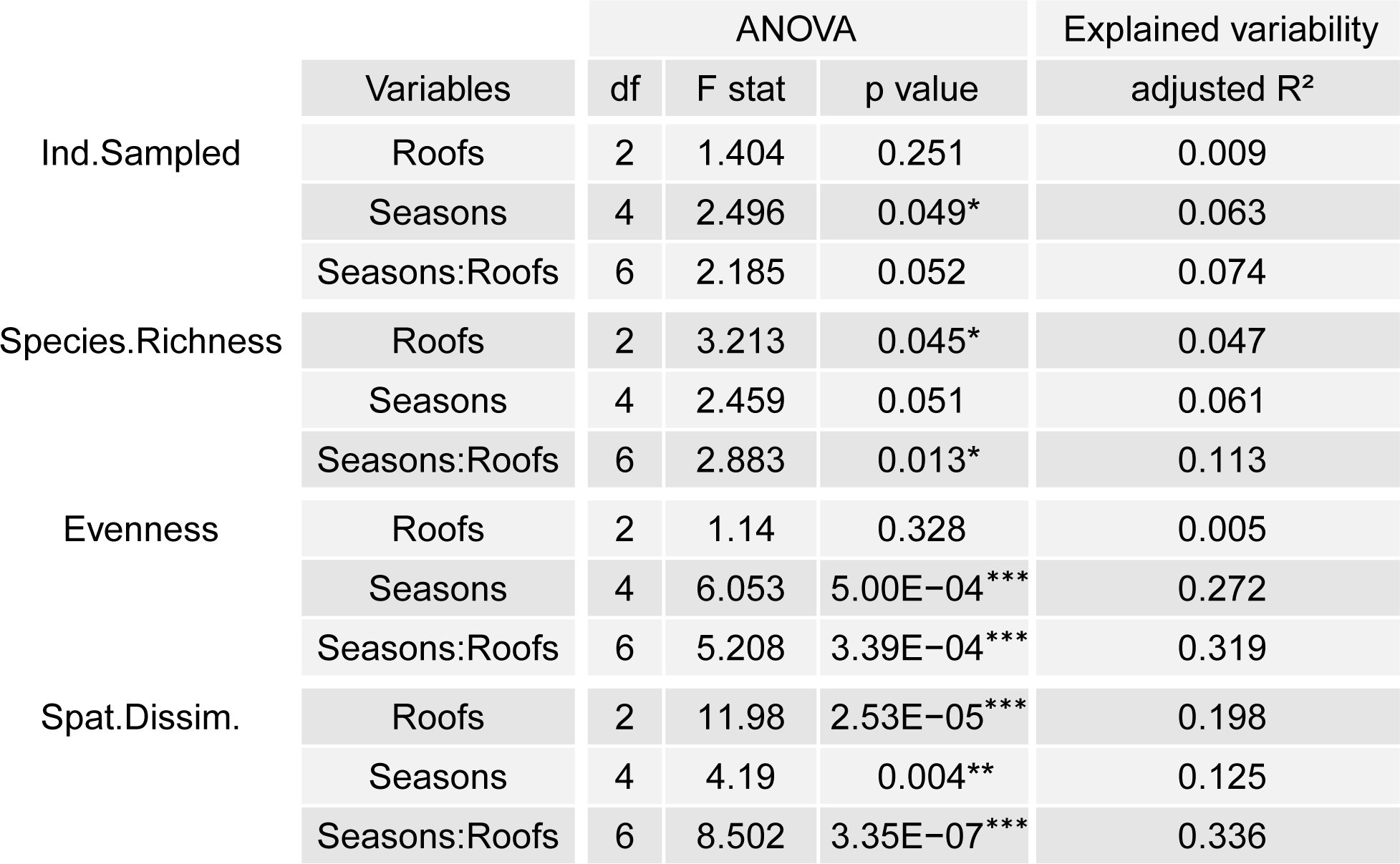
Results of ANOVA analyzing the responses of each community to the roof location, the season and their interaction. The metrics significantly affected by one parameter are indicated by stars (* p-value <0.05; ** p-value <0.01; *** p-value <0.001). The adjusted R^2^ indicates the fraction of the explained variability calculated from a linear model for each variable.

### Modelling the spatio-temporal dynamics of *Adineta sp*

Our simulations best fitted the experimental data when scenario 3 and 5 were run under permissive conditions (Figure 6a). In both scenarios, distinct immigration probabilities among cryptic *Adineta* species were hypothesized suggesting that the dispersal parameter could be the most important one shaping the bdelloid community dynamics. Second, the likelihood of our simulations considerably improved when exponential and normal distributions were used (Figure 6a). On the one hand, the exponential distribution produced communities with one highly dominant species representing 55.1 (for scenario 3) to 73.86% (scenario 5) of the individuals (Figure 6b), as observed experimentally (67%, Figure 7a). On the other hand, more species (up to fourteen) were retrieved over the hundred communities simulated when the normal distribution was used (Figure 6c), best fitting the field data (which retrieved 24 distinct *A. vaga* species). As a result, the combination of those two types of distribution, as used under the hybrid model, produced results that slightly better fitted the experimental data meaning that one species, adapted to the local conditions, outcompetes a few other species that are able to persist in sub-optimal conditions (Figure 6a). Third, the different values of intra and inter-specific competition strengths tested did not affect the model under the combination of parameters tested and was not used in the final simulations using the hybrid model.

Our statistical analysis demonstrated a significant effect of season, with larger and more diversified communities in autumn than in winter. Therefore, we compared the simulations to the communities observed experimentally for each season separately. This suggested that harsher conditions model the communities sampled in winter best. Indeed, the winter communities were better fitted by the harsh and intermediate conditions whatever the scenario (mean = -2.20). In contrast, permissive conditions suit communities from the autumnal, spring and summer sampling best (with pseudolikelihoods mean for autumn and spring = -2.83; for summer mean = -1.62, see Supplemental figure 8).

The Mantel test computed on the entire dataset indicated that 26.88% of the community dissimilarities could significantly be explained by the geographic distance separating the communities (permutations = 1000; p-value = 0.001) suggesting that communities sampled on the same roof were more related than communities of distinct roofs, as observed on Figure 4.

## Discussion

### Dispersal and colonization of experimental habitat patches

We confronted model simulations to a quantitative field experiment to examine the spatial and temporal dynamics of bdelloid rotifer community composition. Our model fitted the data best when A. vaga species differed in their immigration probability, but only when this probability covaried positively with survival and reproduction probability (scenario 5), or when all species had the same survival and reproduction probability (scenario 3). Those results highlight that immigration is an important parameter in bdelloid dynamics while the effect of reproduction and survival (*i.e.* fitness) was low but not negligible, at least within the range of values tested. Even though our simulations highlight the importance of dispersal in bdelloid community dynamics, it is the order of magnitude in the differences among species that fitted the experimental data best. Indeed, switching from a linear distribution of immigration probability to the hybrid between the exponential and normal distributions increased the pseudolikelihood of the model considerably for scenario 3 and 5. This observation does not automatically imply that *Adineta* species have distinct dispersal capacities but rather that immigration varies between species. If all species have similar dispersal capacities, as could be expected here since the sampled *A. vaga* individuals have similar morphological characteristics, the chances to immigrate may be positively correlated with its presence in the nearby metacommunity (source-sink dynamic). This is corroborated by our results showing that the most abundant *A. vaga* species in our study (species 9 representing 67.41% of all individuals genotyped during two years, Figure 4) was omnipresent in Belgium and especially in the park in Namur (Figure 3) confirming that immigration from the local meta-community was the most critical parameter under the conditions tested.

The spatial β-diversity calculated from our simulations suggested that most communities are similar at early stages of colonization, indicating that the same species tend to arrive first in most dishes (Figure 7). As a consequence, most *A. vaga* species appear to have low chances to colonize our experimental dishes, a few species have fair chances and only a restricted number of *A. vaga* species have probabilities high enough to immigrate in multiple dishes within the time window of the study. The species with higher chances to immigrate correspond to the species present in the same geographical region, being here *A. vaga* species 9 who is abundant in the nearby park and our experimental dishes. Species with fair chances to immigrate may be the ones from more distant areas (even other European countries) as three species were found across the continent (species 11, 12 and 14, Supplemental figure 5). Finally, the two species (3 and 5) found in our communities that were also sampled in the US (Supplemental figure 5) could be indicative of low, but nonzero, immigration probabilities for geographically distant bdelloid species as previously highlighted by Fontaneto *et al* (2007). After colonization, species with the highest survival and reproduction probabilities started to expand while some new species immigrated independently in the different communities (the spatial β-diversity slowly decreased until week 12, Figure 7b). However, those new species did not develop much as the median β-diversity stayed above 0.7 (Figure 7b), indicating that the communities present in the distinct Petri dishes were similar, with only a few species unique to any particular dish. This result highlights that only few *A. vaga* species immigrated and started reproducing upon settlement, while rare species arrived and survived in suboptimal conditions but did not reproduce implying that less suitable genotypes become displaced by the best one.

However, rarer immigrations in specific communities, such as on roof A where the diversity was lower (richness, Table 1) and where species 9 was less retrieved (60% petri-dishes without species 9 on roof A), may have left available space for other *A. vaga* species from more distant areas to settle (such as species 10, 13-18, Figure 3 and 4). We do not know which parameter did limit the dispersal on roof A by *A. vaga* species 9, but this roof was surrounded by buildings while C and B were more isolated and closer to the park in which species 9 is extremely abundant. Moreover, Fontaneto *et al* (2008) showed that although *A. vaga* geographical distribution was negatively correlated with species delimitation resolution (*i.e.* bdelloid morphospecies are cosmopolitan whereas genetic clusters are generally locally distributed), some cases of large distribution were detected for the lowest taxonomic rank as observed here for species 3, 5, 11, 12 and 14 that had already been isolated in other European countries or even in the US (Supplemental figure 5).

From a theoretical point of view, scenario 2 where fitness varies across species but dispersal is identical, may produce a similar community structure with all species able to immigrate but only a few surviving in the newly colonized patch. Yet, simulations under scenario 2 produced communities with a lower pseudolikelihood (mean = -2.7 and -4.3 for the exponential and normal distribution respectively) fitting our field data when compared to scenario 3 (mean pseudolikelihood = -2.3 and -2.8 for the exponential and normal distribution respectively) and scenario 5 (mean pseudolikelihood = -2.2 and -2.8 for the exponential and normal distribution respectively). One possibility is that the immigration rate was too low for all *A. vaga* species in scenario 2 under the tested values; increasing the immigration rates four-fold indeed slightly improved the model fit for scenario 2 (mean pseudolikelihood = -2.5 and -3.1 for the exponential and normal distribution, data not shown). However, only the species evenness and β-diversity generated a better fit. The species richness and abundance of individuals did not fit with the experimental data: per patch 20 species representing a mean of 80 individuals were retrieved under these simulated parameters of scenario 2, whereas field data found per patch 1-5 species representing a mean of 16.5 individuals.

Thus, efficient immigration from the metacommunity combined with effective survival and reproduction (scenario 5) would explain the diversity of cryptic *Adineta* species observed here and principally the dominance of *A. vaga* species 9 on roofs C and B. The lower richness observed on roof A could be the consequence of a more limited immigration on roof A because roofs C and B were located at the margins of Park Louise-Marie whereas roof A is located in-between buildings and less exposed to wind.

### Community development throughout seasons

Another important result inferred from our model is the impact of seasons on the dynamics of bdelloid rotifer communities. When parameterized for harsh and intermediate conditions, the model predictions fitted the winter samples best. Spring, summer and autumn samples were best predicted by the permissive conditions. The main factor distinguishing winter from other seasons was lower daily temperatures (Supplemental figure 6) with frequent periods of frost. In another study on the temporal dynamics of *Macrotrachela quadricornifera* populations from mosses, Ricci *et al* (1989) concluded that the number of isolated individuals correlated with the average relative humidity prior sampling but not with temperature or rainfall. Indeed, relative humidity of the air impacts bdelloid community dynamics since population growth is arrested during periods of desiccation while reproduction is boosted immediately after rehydration (Ricci *et al*, 2007). Nevertheless, temperature also plays a role on their reproductive rate as observed in our study. Indeed, six out of the nine petri-dishes sampled on the roofs in which no *Adineta* individual were found (7 trimestral dishes and 2 yearly dishes, Supplemental table 1 and Figure 7a), were retrieved in winter and abundance was significantly lower during this season (Figure 5). We also did not observe any other bdelloid families in those six winter dishes (pers. obs.). The model suggested that some individuals arrived in those communities but that conditions were too harsh for any bdelloid species to persist over several time-steps (Supplemental figure 7). It is unlikely that those dishes were empty due to a less efficient passive dispersal in winter as the corresponding two-yearly dishes, sampled in winter, were also empty. The yearly dishes remained exposed to dispersal by wind for an entire year and it is therefore unlikely that no individuals ever colonized those dishes when all dishes sampled in summer or autumn were inhabited. The empty yearly dishes we retrieved reveal strong effects of winter conditions on *A. vaga* communities. Temperatures were indeed significantly lower in winter, especially between December and January (Supplemental table 4), hindering population growth and explaining, at least partially, the significantly lower abundance retrieved during winter. In contrast, the higher abundance observed in autumnal communities (Figure 5) was correlated with higher humidity (Supplemental figure 6) which enables rehydration and facilitates population growth from dried bdelloid rotifer propagules spread by the wind throughout the year and reproducing asexually. No significant abundance differences were observed between spring and summer, which were similar in terms of meteorological conditions, except for the higher temperatures observed during summer that may be beneficial for population growth to a certain extent.

**Figure 5:**
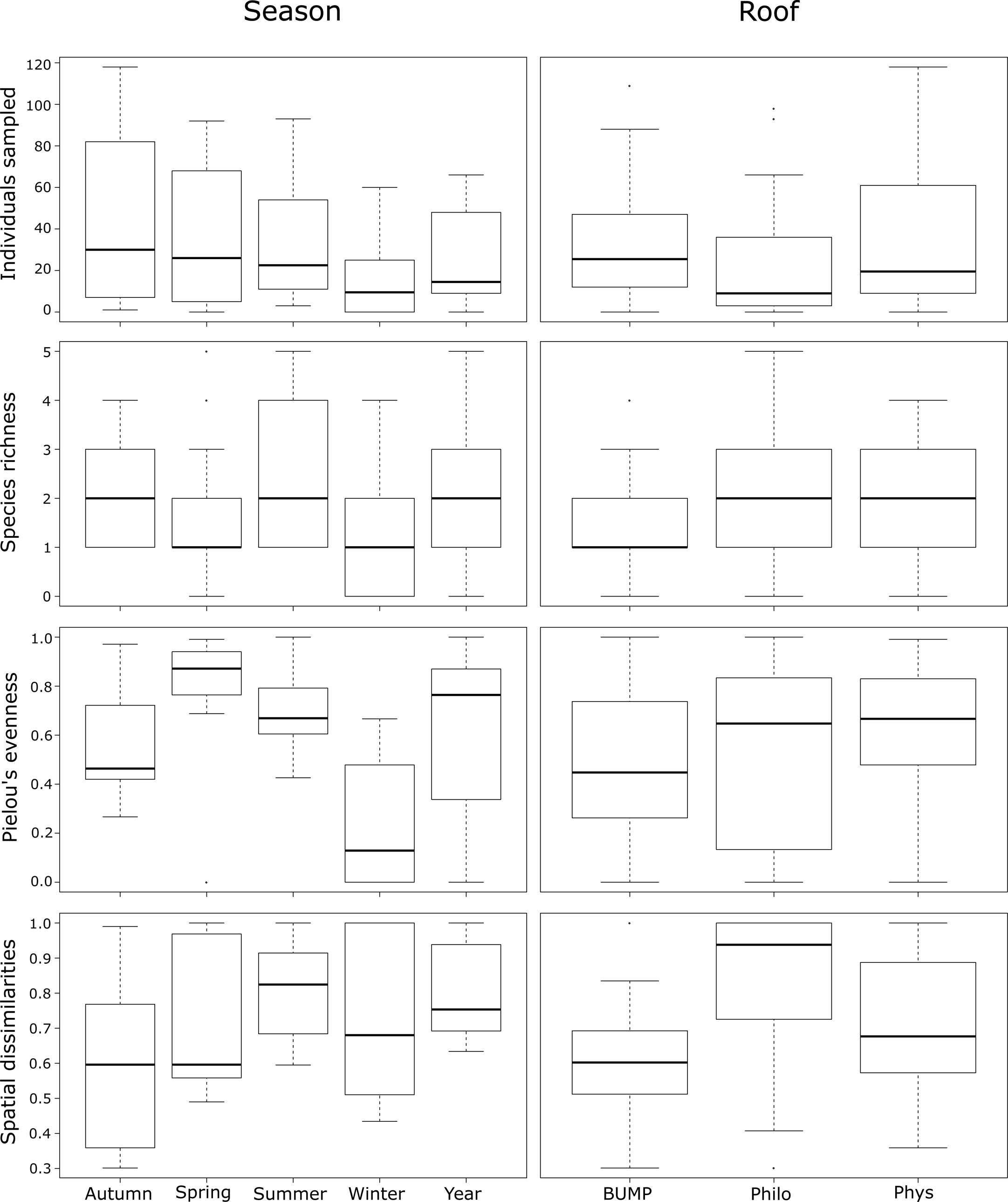
Summary statistics boxplots by seasons and roofs. The number of individuals sampled, the species richness, the Pielou’s index of species evenness and the spatial dissimilarities are presented among communities grouped by seasons and roofs.

**Figure 6:**
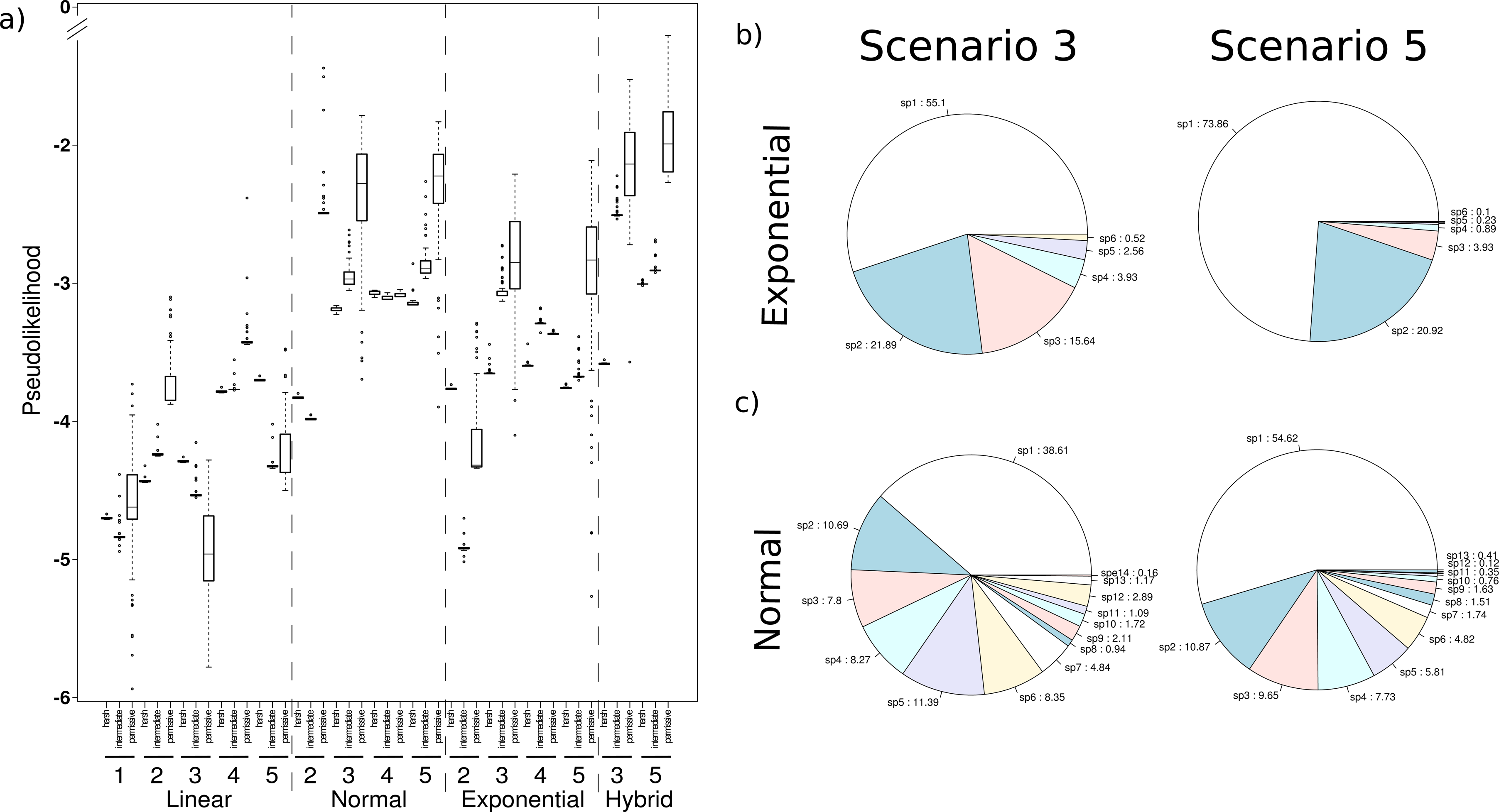
Model fit to the experimental data. a) Likelihoods of the simulated communities fitting the roof data under linear, exponential (only scenario 2-5), normal (only scenario 2-5) and hybrid distributions (only scenario 3 and 5) for the probabilities of immigration, survival and reproduction at each time-step. Pseudolikelihood values closer to 0 indicate better fit of the simulations to the data collected from the roofs experiment. b) and c) are pie charts representing the proportion of each species in the communities simulated by the model according to exponential and normal distributions. The results presented are the simulation of scenario 3 and 5 under permissive conditions which gave the highest likelihood to fit the experimental data from the roofs.

Using the scenarios that best fitted the data (3 and 5 under the hybrid distribution), we used the model to simulate the history of the communities sampled on our three roofs, as only final composition was observed empirically. These simulations suggest that most of the petri-dishes were rapidly colonized, as 25.7 and 23.4% (scenario 3 and 5, respectively) of the communities already harbored at least one individual after the first time-step (Figure 7a). Following colonization, most communities were growing exponentially until week 12, with 93.6 and 84% of the plates being colonized for scenario 3 and 5 respectively (experimentally we observed that 90% of the plates were colonized after two years). In general, the species with the highest immigration probability dominated the community and represented 40.23 and 76.15% of all the individuals sampled according to scenario 3 and 5, respectively (Figure 7a). Yet, new species still arrived and were able to persist even though the primary colonizer had already settled (abundance exponentially increasing and evenness reaching a plateau, Figure 7b). The simulated β-diversity dynamics suggest that, at the onset, communities were quite similar, with the same species immigrating in the different Petri dishes (Figure 7b). As also rarer species immigrated over time, β-diversity slowly decreased. At the end of the simulation, only 6.4% (scenario 3) and 16.0% (scenario 5) of the dishes were empty (Figure 7a) (experimentally we observed that 10% of the plates were empty after two years). Some of those empty dishes were colonized during earlier time-steps but the immigrating individuals could not reproduce or died rapidly, while other dishes were never colonized (Supplemental figure 7).

**Figure 7:**
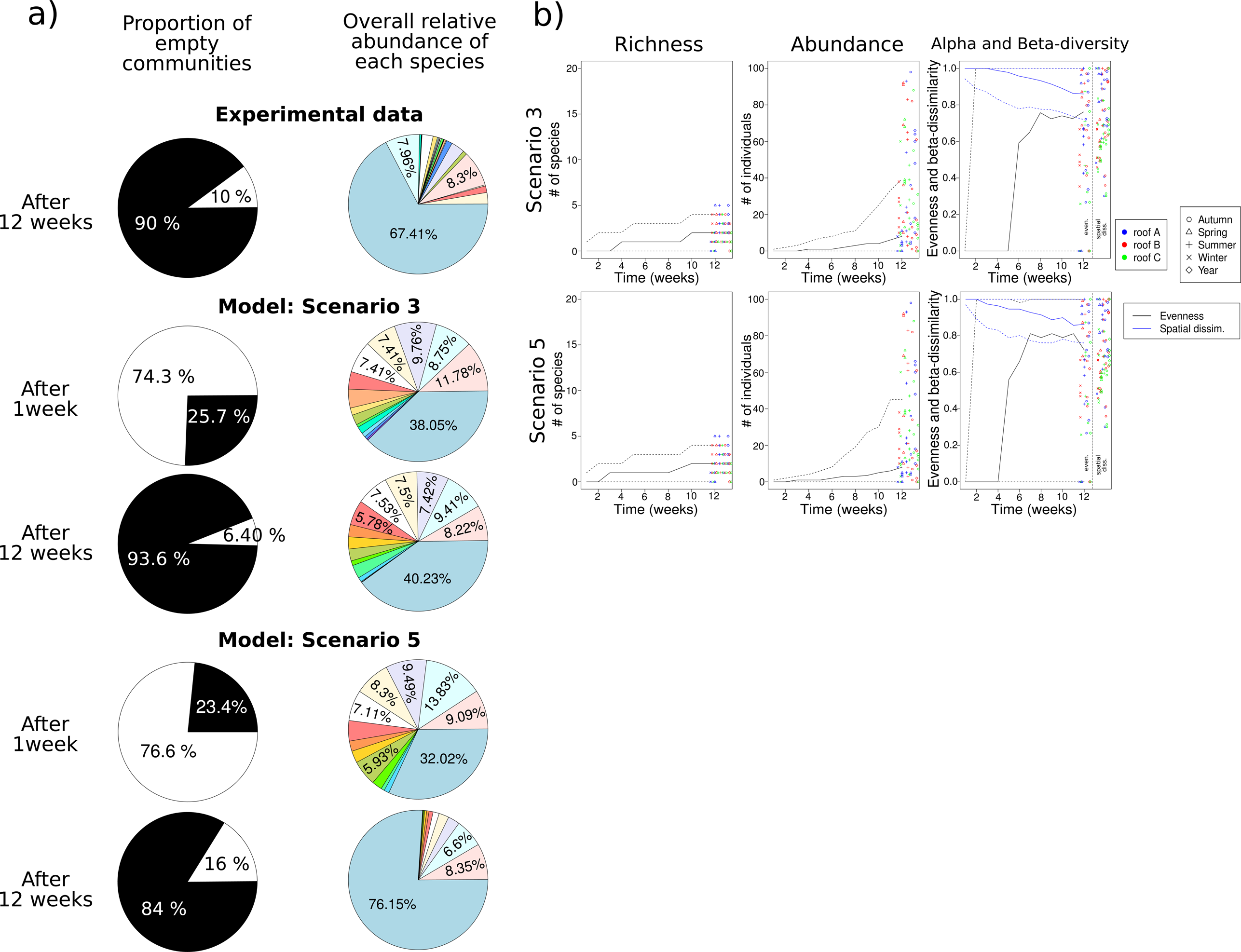
Fit of the Adineta vaga communities simulated according to the hybrid model with the experimental data. a) The overall number of colonized roof communities at sampling time (12 weeks) was 90% in the experimental dataset, while it reached 23.4-25.7% after 1 week or 84-93.6% after 12 weeks (i.e. time-steps) for the simulated communities. The roof communities were colonized by 24 species in the experiment and the relative abundance (not shown when <5%) of the dominant species represented 67.41%. Under scenario 3, simulated communities contained 20 species after 12 weeks, one of which accounted for 40.23% of individuals whereas 14 species dominated by one species (76.15%) were present in the simulations under scenario 5. b) The quantiles (Q50 plane line; Q05 and Q95, dotted lines) for species richness, abundance of individuals and diversity (evenness and spatial dissimilarities) were calculated at each time-step (12 weeks) for the simulated community (100 simulations) and plotted with the results of the roofs data (colored symbols) for comparison.

Several studies on monogonont rotifers, a sister clade of bdelloids, described an increase in generation time (due to lower reproductive rate and longer lifespan) when temperature decreased from 30°C to 15°C (Xiang *et al*, 2010; Kauler and Enesco, 2011). The same applies to the bdelloid rotifer *A. vaga* for which the number of laid eggs was lower and the egg development time was longer when individuals and eggs were incubated at 4°C instead of 25°C. Indeed less than 14% of the females laid one egg after 6 days at 4°C while at 25°C 87.5% of the females laid eggs within the first 20h, and 0 eggs hatched after 13 days at 4°C versus 40% hatching after 48h at 25°C (M. Terwagne and L. Herter, personal communication). Strikingly, all the eggs incubated at 4°C for 13 days rapidly hatched when put back at 25°C. Thus, freezing periods combined with low temperatures and low reproductive rates, will tend to reduce population sizes in winter in the bdelloid rotifer species *A. vaga*, followed by population growths when temperature increases. Extremely high temperatures, may result in fully desiccated communities that entered in a paused metabolic state, continuing reproduction upon rehydration. Those observations suggest that although A. vaga communities could survive throughout the year, the harsher conditions encountered in winter may result in an annual bottleneck followed by expansion of the few remaining individuals reaching a climax in autumn when temperatures and humidity are more stable. A second hypothesis would be that in autumn the conditions were more suitable for dispersal with a continuous immigration of the species from the metacommunity. However, this is less likely as, in our model, immigration did not co-vary with harshness (Supplemental figure 3) and the maximum wind speed was not higher in autumn (Supp. Fig. 6). This suggests that the main parameters influencing population growth throughout the year are survival and reproduction. Moreover, in such harsh environments where mortality is high it is advantageous to be asexual since one surviving individual can re-start the population. It is therefore not surprising to encounter in ephemeral habitats, such as our experimental petri-dishes, mostly bdelloid rotifers (pers. obs.).

### Co-existence and differential adaptation among cryptic species

*Adineta* species 8 and 13 were also frequent and abundant throughout this study (93 and 97 individuals genotyped in 22 and 18 roof patches respectively; Figure 4) while species 13 was never sampled in the park next to University and species 8 was never detected before. The remaining 21 *Adineta* species identified in our petri-dishes appeared sporadically (less than ten times within the roof patches, with seven species only found once, Figure 4). A similar observation was made with *M. quadricornifera* populations where one electromorph was retrieved several times within a moss throughout their two-year study, representing 78.7% of the community, while the four other electromorphs appeared occasionally (Ricci et al, 1989). These results suggest that distinct cryptic species co-occur periodically but that only few clones expand significantly or immigrate and settle frequently as depicted by our simulations under scenario 5 and 3 respectively (Figure 1 and Supplemental figure 3).

We also observed that eight species (species 13 to 18, 22 and 24), also present in Belgium but not in Namur, were almost exclusively found on roof A, where the dominant species 9 was less abundant (Figure 4). This may reveal local adaptation of those eight species to environmental conditions specific to roof A and/or limited dispersal of species 9 to this roof providing space for other species to settle. This may also suggest that species 9 presents a higher competitive strength than species 13 to 18, 22 and 24 which can only settle and expand when species 9 is absent, or rare (Figure 3 and 4). This remains speculative since we could not find any differences in the community modelled by the three degrees of intra and inter-species interaction strength. A similar case, although rare, was observed for species 4 which was only detected two times in summer (12 and 18 individuals), but both times species 9 was absent. In this latter case, dispersal cannot explain the absence of the dominant species 9 as it was present in the other replicates from the same roof. The dominant abundance of species 4 in those two dishes could highlight adaptation to particularly dry and warm conditions or a difference in desiccation tolerance among cryptic *Adineta* species.

To our knowledge, no studies have focused on habitat specialization, neither on desiccation tolerance differences among cryptic species of bdelloid rotifers. In the first case, it seems tricky to distinguish between the presence of a species due to habitat specialization or to its high abundance in the metacommunity acting as a permanent source. In the second case, several reports show that there are several degrees of tolerance to desiccation among bdelloid morphospecies (Eyres et al, 2015) but none have focused on variations within species complexes. However, three cryptic species morphologically identified as *Rotaria rotatoria* have been reported to have distinct temperature preferences (Xiang *et al*, 2016), often linked to desiccation tolerance. In the present study, we did not identify physico-chemical parameters that may vary among the different dishes. However, if there are differences between dishes, they are probably due to variations between roofs rather than differences among dishes as the three replicates of each roof tended to give similar results. In addition, the meteorological records that we used were not precise enough to make any distinction between roofs although we doubt there were significant climatic differences between dishes located < 150 meters apart. The differences between roofs are more likely resulting from the community assemblage itself. We did not identify the number of other zooplankton and phytoplankton species present in the dishes but we observed that roof A communities were, in general, poor whereas C and B communities often presented several bdelloid species and a few other animals being desiccation tolerant such as tardigrades and nematodes while being crowded with algae (pers. obs.). It is not surprising to find those taxa in our communities as their dispersal capacity is comparable to rotifers (tardigrades and nematodes can also desiccate at any stage in their life-cycle) and they are often transported together on sediments by the wind (Nkem *et al*, 2006).

Interestingly, we did not retrieve the A. vaga species that was exclusively present in the soil patches around Namur (species E in Debortoli et al, 2016), showing that bdelloid species dwelling in lichen patches on trees more exposed to wind have a higher chance to disperse than species from soil habitats, frequently encountering parasites (Wilson et al, 2011).

## Conclusion

Our results simulating the dynamics of bdelloid rotifer communities showed that ecological scenarios in which cryptic *Adineta* species present distinct immigration probabilities fitted the experimental data more accurately. This observation highlights the prevalent role of passive dispersal in the dynamics shaping bdelloid rotifer communities. Our model emphasized the distinct dispersal probabilities among *A. vaga* species with the most abundant *A. vaga* species in our study being the one present in the nearby area (Species A, Debortoli et al. 2016), suggesting that subsequent short-range dispersals from the metacommunity are more frequent than long-range dispersals. Combined with their desiccation tolerance and asexual mode of reproduction, dried bdelloid individuals can start new populations upon hydration and multiply, as observed here during summer and autumn conditions. Moreover, asexual bdelloid populations experiencing strong bottlenecks (in winter periods) easily re-established the population. In harsh, ephemeral environments where survival is hard and driven by density-independent factors, asexual reproduction is an advantage compared to sexual reproduction because of its higher productivity (Scheu & Drossel, 2007; Song et al., 2012). As a consequence, the bdelloid rotifers, the most notorious asexual animal clade, could persist in the long term through spatio-temporal dynamics in which abundant clones easily disperse as desiccated tuns and settle in suitable habitats, even harsh ones where the death rate is high and where parasites are absent (Wilson et al., 2013), enabling their long-term persistence as asexuals.

## Author contributions

Design of the study: N. D. and K. V. D.

Rotifer sampling, DNA preparation and genetic analyses: N. D.

Community structure analyses: N. D.

Model development: N. D. and F. D. L.

Core writing of the manuscript: N. D., F. D. L. and K. V. D.

## References

Amarasekare, P., & Nisbet, R. M. (2001). Spatial heterogeneity, source-sink dynamics, and the local coexistence of competing species. The American Naturalist, 158(6), 572–584.

Bass, D., & Cavalier-Smith, T. (2004). Phylum-specific environmental DNA analysis reveals remarkably high global biodiversity of Cercozoa (Protozoa). International Journal of Systematic and Evolutionary Microbiology, 54(6), 2393–2404.

Bray, J. R., & Curtis, J. T. (1957). An ordination of the upland forest communities of southern Wisconsin. Ecological Monographs, 27(4), 325–349.

Costello, M. J., & Chaudhary, C. (2017). Marine Biodiversity, Biogeography, Deep-Sea Gradients, and Conservation. Current Biology, 27(11), R511–R527.

De Laender, F., Melian, C. J., Bindler, R., den Brink, P. J., Daam, M., Roussel, H., … & Janssen, C. R. (2014). The contribution of intra-and interspecific tolerance variability to biodiversity changes along toxicity gradients. Ecology letters, 17(1), 72–81.

Debortoli, N., Li, X., Eyres, I., Fontaneto, D., Hespeels, B., Tang, C. Q., … & Van Doninck, K. (2016). Genetic exchange among bdelloid rotifers is more likely due to horizontal gene transfer than to meiotic sex. Current Biology, 26(6), 723–732.

Donner, J. (1965). Ordnung Bdelloidea. Bestimmungsbücher zur Bodenfauna Europas 6: 1–297.

Drummond, A. J., & Rambaut, A. (2007). BEAST: Bayesian evolutionary analysis by sampling trees. BMC evolutionary biology, 7(1), 214.

Eyres, I., Boschetti, C., Crisp, A., Smith, T. P., Fontaneto, D., Tunnacliffe, A., & Barraclough, T. G. (2015). Horizontal gene transfer in bdelloid rotifers is ancient, ongoing and more frequent in species from desiccating habitats. BMC biology, 13(1), 90.

Finlay, B. J. (2002). Global dispersal of free-living microbial eukaryote species. Science, 296(5570), 1061–1063.

Flot, J. F., Hespeels, B., Li, X., Noel, B., Arkhipova, I., Danchin, E. G., … & Barbe, V. (2013). Genomic evidence for ameiotic evolution in the bdelloid rotifer Adineta vaga. Nature, 500(7463), 453–457.

Folmer, O., Black, M., Hoeh, W., Lutz, R., & Vrijenhoek, R. (1994). DNA primers for amplification of mitochondrial cytochrome c oxidase subunit I from diverse metazoan invertebrates. Molecular marine biology and biotechnology, 3(5), 294–299.

Fontaneto, D., Ficetola, G. F., Ambrosini, R., & Ricci, C. (2006a). Patterns of diversity in microscopic animals: are they comparable to those in protists or in larger animals?. Global Ecology and Biogeography, 15(2), 153–162.

Fontaneto, D., & Ricci, C. (2006b). Spatial gradients in species diversity of microscopic animals: the case of bdelloid rotifers at high altitude. Journal of Biogeography, 33(7), 1305–1313.

Fontaneto, D., Herniou, E. A., Barraclough, T. G., & Ricci, C. (2007). On the global distribution of microscopic animals: new worldwide data on bdelloid rotifers. ZOOLOGICAL STUDIES, 46(3), 336.

Fontaneto, D., Barraclough, T. G., Chen, K., Ricci, C., & Herniou, E. A. (2008). Molecular evidence for broad-scale distributions in bdelloid rotifers: everything is not everywhere but most things are very widespread. Molecular Ecology, 17(13), 3136–3146.

Fontaneto, D., Westberg, M., & Hortal, J. (2011). Evidence of weak habitat specialisation in microscopic animals. PLoS One, 6(8), e23969.

Fontaneto, D., & Barraclough, T. G. (2015). Do species exist in asexuals? Theory and evidence from bdelloid rotifers. Integrative and comparative biology, 55(2), 253–263.

Fujisawa, T., & Barraclough, T. G. (2013). Delimiting species using single-locus data and the Generalized Mixed Yule Coalescent approach: a revised method and evaluation on simulated data sets. Systematic biology, 62(5), 707–724.

Guidetti, R., & Jönsson, K. I. (2002). Long-term anhydrobiotic survival in semi-terrestrial micrometazoans. Journal of Zoology, 257(2), 181–187.

Hart, S. P., Schreiber, S. J., & Levine, J. M. (2016). How variation between individuals affects species coexistence. Ecology letters, 19(8), 825–838.

Hubbell, S. P. (2001). The Unified Neutral Theory of Biodiversity and Biogeography, vol. 32. Princeton University Press.

Katoh, K., & Standley, D. M. (2013). MAFFT multiple sequence alignment software version 7: improvements in performance and usability. Molecular biology and evolution, 30(4), 772–780.

Kauler, P., & Enesco, H. E. (2011). The effect of temperature on life history parameters and cost of reproduction in the rotifer Brachionus calyciflorus. Journal of Freshwater Ecology, 26(3), 399–408.

Lopes, P. M., Bozelli, R., Bini, L. M., Santangelo, J. M., & Declerck, S. A. (2016). Contributions of airborne dispersal and dormant propagule recruitment to the assembly of rotifer and crustacean zooplankton communities in temporary ponds. Freshwater Biology, 61(5), 658–669.

Lowe, W. H., & McPeek, M. A. (2014). Is dispersal neutral?. Trends in ecology & evolution, 29(8), 444–450.

Maynard Smith, J. (1978). The evolution of sex. CUP Archive.

Merckx, T., Souffreau, C., Kaiser, A., Baardsen, L. F., Backeljau, T., Bonte, D., … & Van Dyck, H. (2018). Body-size shifts in aquatic and terrestrial urban communities. Nature, 558, 113–116.

Nkem, J. N., Wall, D. H., Virginia, R. A., Barrett, J. E., Broos, E. J., Porazinska, D. L., & Adams, B. J. (2006). Wind dispersal of soil invertebrates in the McMurdo Dry Valleys, Antarctica. Polar Biology, 29(4), 346–352.

Oksanen, J., Kindt, R., Legendre, P., O’Hara, B., Stevens, M. H. H., Oksanen, M. J., & Suggests, M. A. S. S. (2007). The vegan package. Community Ecology Package, 10, 631–637.

Pielou, E. C. (1966). Species-diversity and pattern-diversity in the study of ecological succession. Journal of Theoretical Biology, 10(2), 370–383.

Pons, J., Barraclough, T. G., Gomez-Zurita, J., Cardoso, A., Duran, D. P., Hazell, S., … & Vogler, A. P. (2006). Sequence-based species delimitation for the DNA taxonomy of undescribed insects. Systematic biology, 55(4), 595–609.

Ricci, C. (1983). Life histories of some species of Rotifera Bdelloidea. Hydrobiologia, 104(1), 175–180.

Ricci, C., Pagani, M., & Bolzern, A. M. (1989). Temporal analysis of clonal structure in a moss bdelloid population. In Rotifer Symposium V (pp.145–152). Springer Netherlands.

Ricci, C. (1998). Anhydrobiotic capabilities of bdelloid rotifers. Hydrobiologia, 387, 321–326.

Ricci, C., & Caprioli, M. (2005). Anhydrobiosis in bdelloid species, populations and individuals. Integrative and Comparative Biology, 45(5), 759–763.

Ricci, C., Caprioli, M., & Fontaneto, D. (2007). Stress and fitness in parthenogens: is dormancy a key feature for bdelloid rotifers?. BMC Evolutionary Biology, 7(2), S9.

Soininen, J., McDonald, R., & Hillebrand, H. (2007). The distance decay of similarity in ecological communities. Ecography, 30(1), 3–12.

Tamura, K., & Nei, M. (1993). Estimation of the number of nucleotide substitutions in the control region of mitochondrial DNA in humans and chimpanzees. Molecular Biology and Evolution, 10(3), 512–526.

Tamura, K., Peterson, D., Peterson, N., Stecher, G., Nei, M., & Kumar, S. (2011). MEGA5: molecular evolutionary genetics analysis using maximum likelihood, evolutionary distance, and maximum parsimony methods. Molecular biology and evolution, 28(10), 2731–2739.

Tang, C. Q., Humphreys, A. M., Fontaneto, D., & Barraclough, T. G. (2014). Effects of phylogenetic reconstruction method on the robustness of species delimitation using single-locus data. Methods in Ecology and Evolution, 5(10), 1086–1094.

Wilson, C. G., & Sherman, P. W. (2010). Anciently asexual bdelloid rotifers escape lethal fungal parasites by drying up and blowing away. Science, 327(5965), 574–576.

Wilson, C. G. (2011). Desiccation-tolerance in bdelloid rotifers facilitates spatiotemporal escape from multiple species of parasitic fungi. Biological journal of the Linnean Society, 104(3), 564–574.

Wilson, C. G., & Sherman, P. W. (2013). Spatial and temporal escape from fungal parasitism in natural communities of anciently asexual bdelloid rotifers. Proceedings of the Royal Society of London B: Biological Sciences, 280(1765), 20131255.

Xiang, X. L., Xi, Y. L., Zhang, J. Y., Ma, Q., & Wen, X. L. (2010). Effects of temperature on survival, reproduction, and morphotype in offspring of two Brachionus calyciflorus (Rotifera) morphotypes. Journal of Freshwater Ecology, 25(1), 9–18.

Xiang, X., Jiang, R., Tao, Y., Chen, Y., & Xi, Y. (2016). Differences in life history characteristics among three sympatric evolutionary species of the Rotaria rotatoria complex. Journal of Freshwater Ecology, 31(3), 351–360.

Zhao, S., Guo, Y., Sheng, Q., & Shyr, Y. (2014). Heatmap3: an improved heatmap package with more powerful and convenient features. BMC Bioinformatics, 15(S10), P16.

